# Modeling metabolic variation with single-cell expression data

**DOI:** 10.1101/2020.01.28.923680

**Authors:** Yuanchao Zhang, Man S. Kim, Elizabeth Nguyen, Deanne M. Taylor

**Affiliations:** Department of Biomedical and Health Informatics, The Children’s Hospital of Philadelphia, Philadelphia, PA 19041, USA; Department of Genetics, Rutgers University, Piscataway, NJ 08854, USA; Department of Pediatrics, Perelman School of Medicine, University of Pennsylvania, Philadelphia, Pennsylvania, 19104, USA

## Abstract

Cellular metabolism encompasses the biochemical reactions and transportation of various metabolites in cells and their surroundings, which are integrated at all levels of cellular functions. We developed a method to systematically simulate cellular metabolism using single-cell RNA-seq (scRNA-seq) data through constraint-based context specific metabolic modeling. We simulated the NAD^+^ biosynthesis activity in 7 different mouse tissues, and the simulated NAD^+^ biosynthesis flux levels showed significant linear correlation with experimental measurements in previous research. We also show that the simulated NAD^+^ biosynthesis fluxes are reproducible using two additional scRNA-seq datasets.

## Introduction

Understanding metabolic strategies used by cells in health and disease could help contrast and identify key network points for therapeutic development. For example, many cancer cells have high levels of glucose consumption and lactate secretion called the Warburg effect (Potter et al., 2016; Warburg et al., 1927) and manipulating metabolic pathways has been shown to overcome tumor immune evasion (Leone et al., 2019). Inborn errors of metabolism (IEMs) comprise the largest category of inheritable human diseases, which include over 500 diseases caused by mutations in genes that have functions in metabolism (Childs et al., 2001; DeBerardinis and Thompson, 2012) that often lead to devastating and complex development comorbidities, including neurodegeneration and physical incapacitation (Pierre, 2013).

Roadblocks to diagnosing and developing therapeutics for many metabolic disorders result from a lack of understanding of the origins of their heterogeneous clinical phenotypes. The heterogeneity is likely due to the manifestation of a combination of metabolic dysfunctions in various cell and tissue types. Modeling cell- and tissue-type specific metabolism would therefore be useful for generating hypotheses on how cell types may utilize different metabolic strategies in an *in vivo* context. Towards this purpose, we have both developed new methods and adapted existing methods to systematically create *in silico* metabolic models of cell and tissue types from single-cell RNA-seq data.

Analysis of single-cell RNA-seq data allows for the definition of “cell type” based on shared characteristics in single-cell transcriptomic profiles. This type of high-resolution transcriptomic characterization of cells and tissues offers significant potential in understanding the function of cells and tissues, including cell- and tissue type specific metabolic processes that can be inferred through context specific metabolic modeling (Pacheco et al., 2015). Metabolism can be modeled as a collection of metabolite conversions within a cellular compartment and exchanges between different subcellular compartments, such as the cytoplasm, nucleus, and mitochondrial matrix (**Figure S1A** and **S1B**). Comprehensive metabolic network models of many organisms are curated by other groups and provided as community resources (King et al., 2016). We can select a subset of metabolic reactions from a metabolic network model of interest to represent cell- or tissue-type specific metabolism, based on transcription levels of relevant enzymes and transporters from a specific experiment. The selected metabolic reactions then comprise a context specific metabolic model, which can be used to simulate cell- or tissue-type specific metabolic fluxes through various computational methods (Bordbar et al., 2014), such as flux balance analysis (FBA) (Orth et al., 2010), flux variability analysis (FVA) (Mahadevan and Schilling, 2003), and uniform flux sampling (Schellenberger and Palsson, 2009). The simulated metabolic fluxes can be interpreted in biological contexts to hypothesize the metabolic variability between different cell- and tissue-types, and the hypotheses can further be tested by experimental profilings of metabolism.

We applied constraint based metabolic modeling method on several mouse scRNA-seq datasets to simulate the metabolic fluxes of various cell and tissue types. We constructed context specific metabolic models for each cell and tissue type in the scRNA-seq datasets using CORDA (Schultz and Qutub, 2016) and comprehensive mouse metabolic model iMM1415 (Sigurdsson et al., 2010). The simulated NAD^+^ biosynthesis fluxes of 7 different mouse tissues show significant linear correlation with experimental measurements (Mori et al., 2014), in two different approaches of model construction using the Tabula Muris scRNA-seq dataset (Schaum et al., 2018). In one approach, we constructed models using the cells in each tissue type and directly simulated the tissue-type specific metabolic fluxes. In the other approach, we constructed models using the cells in each cell type, and we used these cell-type specific models to simulate cell-type specific metabolic fluxes. Then, we computed the tissue-specific metabolic fluxes as the arithmetic mean of comprising cell-type specific metabolic fluxes weighted by the number of cells in each cell type. In addition, we also evaluated the variance of the simulated fluxes by bootstrapping the cells in each cell type or tissue, and the flux mean and standard deviation of the bootstrapping samples are able to illustrate the metabolic variations within each group of cells and between groups of cells. We also applied the developed metabolic modeling method on two Allen Brain Mouse Atlas scRNA-seq datasets, and the simulated NAD^+^ biosynthesis fluxes of 11 mouse brain cell types are reproducible when computed using different scRNA-seq datasets.

## Methods

### Single-cell data processing

We obtained the Tabula Muris mouse single-cell RNA-seq count matrix dataset from https://tabula-muris.ds.czbiohub.org (Schaum et al., 2018). The version 8 dataset of the Smart-Seq2 sequencing results of the fluorescence-activated cell sorting (FACS) captured cells was used to construct context specific metabolic models (see the following Methods section for details).

RNA-seq read count matrices from the Allen Brain Atlas mouse data from the visual cortex (VISp) and anterior lateral motor cortex (ALM) datasets were obtained from the Allen Brain Map Portal, located at https://portal.brain-map.org/atlases-and-data/rnaseq. These two datasets contain SMART-Seq4 sequencing results of FACS captured single cells, which were used to construct context specific metabolic models.

In these single-cell RNA-seq datasets, cells are able to be partitioned by labeled cell type, region, or organ. Cell-to-cell normalization was not applied, because the rankings of gene expression levels, rather than the specific values, were used for metabolic modeling. Filtering and further steps taken with the count matrices are discussed below.

### Construction of a context specific metabolic model

We construct context specific metabolic models by extracting a subset of reactions from a reference genome scale metabolic model based on the single-cell gene transcription levels. A reference model contains all known metabolic reactions that can occur in an organism, and a context specific model only contains the metabolic reactions in a specific tissue or cell type. The reactions in a context specific model comprise a subset of the reactions in the reference model, and the subset is selected according to the transcription levels of enzymes in a specific tissue or cell type. Therefore, a context specific metabolic model more accurately represents the metabolism of its corresponding cell type or tissue.

Context specific metabolic models can represent the transcriptome of any group of cells, which is not limited to cell type, tissue, or organ. Although the cellular organisational levels are different, the model abstracts an open chemical system that is able to import metabolites from external sinks, convert the metabolites internally for various biological objectives, and export metabolites to external demands (**Figure S1A**). The sinks of metabolites can be interpreted as large pools of metabolites that are outside the chemical system defined by the metabolic model, and the sink reactions are able to import the metabolites from the external pools to the system. The demands of metabolites can be interpreted as external biochemical components that are able to consume the metabolites, e.g. histone deacetylases use NAD^+^ to remove acetyl groups from histones (Reid et al., 2017), and the demand reactions are able to consume the metabolites in the system.

We group the cells in the Tabula Muris dataset (Schaum et al., 2018) by the tissue type and cell ontology class annotations of each cell. Then, we constructed context specific models for the tissue types and cell ontology classes. The tissue type models represent the overall metabolism of the cells in the tissue, and the cell ontology class models represent the metabolism of specific cell types. We removed cell ontology classes with less than 35 single cells in the FACS dataset, in order to ensure that the transcriptomic profiles of ontology classes can be accurately inferred from the transcriptomic profiles of individual cells. The models are clustered on the Jaccard dissimilarities (See Supplementary Methods for details) by hierarchical clustering with complete linkage, which are plotted in **Figure 2A** and **2B** as heatmaps and dendrograms. Similarly, we group the cells in the Allen Brain Atlas datasets by the cell subclasses in the annotation, and we constructed context specific models for the cell subclasses.

We convert the transcriptomic profiles of a selected group of cells into gene expression confidence scores based on the mean read count rankings of each gene. For each group of cells, we rank the genes by their mean read counts. Then, we assign discrete confidence scores to each gene to represent the likelihood that the gene is expressed in that group of cells. For example, if values for over 10,000 genes were found for a particular cell group, we could give a high confidence score of ‘3’ for the top 2000 genes, a medium confidence score of ‘2’ for the following 4000 genes, and a lower confidence score of ‘1’ to the further 2000 genes. We assign the confidence score of 0 to the next 1000 genes, which represent an unknown likelihood of expression. Finally, we assign confidence score of −1 to the last ranked genes, which represent that the genes are unlikely to be expressed in the cell group as a whole. The choice of the numbers of reactions with different confidence scores depends on the dropout rate and the amount of technical variance in the dataset. If the dropout rate and technical variance is high, it is more likely that important enzymatic genes will be ranked low. These parameters are individually evaluated for each dataset in an iterative fashion, to balance the sensitivity and robustness of the method.

The gene expression confidence scores are further converted to metabolic reaction confidence scores based on the associations between reactions and genes in the reference metabolic model. The reference metabolic model we used to construct context specific metabolic model is iMM1415, which is obtained from Biochemical, Genetic and Genomic (BiGG) Models database (King et al., 2016). The iMM1415 model is reconstructed by Sigurdsson *et al* (Sigurdsson et al., 2010), which contains 3,726 reactions and 2,775 metabolites in 8 cellular compartments. In this model, 2,210 reactions are associated with 1,375 genes, and their associations are represented as boolean algebraic operations with genes as operands. For example, the gene association of the reaction “ATP + D-Fructose 6-phosphate <=> ADP + D-Fructose 1,6-bisphosphate”, which is catalyzed by phosphohexokinase, is “56421 or 18641 or 18642”, where the numbers are Entrez Gene IDs. When evaluating the boolean algebraic operations, the result of an “or” operation is the maximum gene confidence score of two operand genes, and the result of an “and” operation is the minimum gene confidence score of two operand genes.

We construct context specific metabolic models based on the reaction confidence scores using the Cost Optimization Reaction Dependency Assessment (CORDA) algorithm (Schultz and Qutub, 2016). We choose CORDA among many other construction algorithms (Pacheco et al., 2015) due to its efficiency and discretization of reaction confidence scores. CORDA uses linear programming to construct context specific metabolic models, so its time complexity is tractable in constructing thousands of models. The discretization of reaction confidence scores greatly improves the robustness of model construction based on scRNA-seq data, by reducing the variances between different cells within a group. To implement the construction process, we use Python programming language version 3.6.8, cobrapy version 0.14.1 (Ebrahim et al., 2013), and the Python implementation of CORDA version 0.4.2 (https://github.com/resendislab/corda), and for this study we utilized the commercial linear programming solver Gurobi Solver version 8.1.

The CORDA constructed models are further evaluated and modified for quality and simulation purposes. This is especially relevant when modeling groups of small numbers of single cells, where because of drop out, the combined expression profiles may not represent the full metabolic space. In order to ensure the validity of the models, another step is to examine if a small number of essential metabolites, such as H_2_O and NAD^+^, are absent in the CORDA generated models, and if not, to add them into the models before further simulation. When simulating the biosynthesis fluxes of certain metabolites, such as NAD^+^ and ATP, we also add in demand reactions that consume the corresponding metabolites in the CORDA generated models.

### Computational analysis of metabolic fluxes

We use flux balance analysis (FBA) (Orth et al., 2010) (**Figure S2**) to simulate the metabolic fluxes of context specific metabolic models. The FBA objectives are defined to maximize the fluxes of demand reactions of certain metabolites, in order to simulate the maximum amount of metabolites that can be synthesized by the model under steady state. The metabolic variations between different context specific models are represented by the differences in FBA simulated fluxes. To implement the FBA, we use Python programming language version 3.6.8, Cobrapy version 0.14.1 (Ebrahim et al., 2013) and a commercial linear programming solver Gurobi Optimizer version 8.1.

The in vivo NAD biosynthesis fluxes in different mouse tissues are obtained from the experimental measurements made by Mori *et al* (Mori et al., 2014). The fluxes of the two final NAD biosynthetic reactions, which are respectively catalyzed by nicotinamide mononucleotide adenylyltransferase (NMNAT) and NAD synthetase (NADS), are used to compare with our computationally simulated fluxes.

We simulated tissue specific NAD^+^ biosynthesis fluxes using the constructed context specific metabolic models. The NAD^+^ biosynthesis flux is simulated by adding a NAD^+^ demand reaction in the compartment of cytosol and applying FBA to optimize the NAD^+^ demand reaction. The NAD^+^ biosynthesis fluxes simulated using Tabula Muris tissue specific metabolic models are directly compared with the empirically measured NAD^+^ biosynthesis fluxes represented by the sum of NADS and NMNAT fluxes (Mori et al., 2014). With regard to the NAD^+^ biosynthesis fluxes simulated using Tabula Muris cell type specific models, we use the arithmetic mean of the cell types within a tissue type weighted by the number of cells to represent the tissue type specific flux, which are compared with the empirically measured NAD^+^ biosynthesis fluxes (Mori et al., 2014). The linear regression between the simulated NAD^+^ fluxes and the empirically measured ones are computed by the linregress function in scipy package version 1.2.1. The p-value is computed using Wald Test with t-distribution, with the null hypothesis that the slope is zero. We also simulated the NAD^+^ biosynthesis fluxes using the cell type specific metabolic models constructed using Allen Brain Atlas datasets, and the simulated fluxes of the same cell types are shown to be reproducible.

### Estimating metabolic variation with bootstrapping

In order to estimate the metabolic variation within a group of cells, we construct context specific models based on the bootstrapping samples of the group of cells, and we simulate metabolic fluxes using the constructed models. For each group of cells, we generate 20 bootstrapping samples by sampling 80% of the cells with replacement, and we construct context specific models based on each sample and simulate the metabolic fluxes using the procedures described above. Then, we use the standard deviation of the simulated fluxes of the bootstrapping samples to describe the metabolic variation within the group.

## Results

### Tissue and cell type specific metabolic models

We constructed context specific metabolic models for 20 tissue types and 111 cell types using the 44,728 filtered and annotated FACS sorted cells that are sequenced with Smart-Seq2 (Schaum et al., 2018). We characterized the differences between different models by the Jaccard dissimilarities of the metabolic reactions, which are illustrated as heatmaps (**Figure 1A** and **1B**). We also illustrated the differences between different models by applying UMAP dimensionality reduction on the Jaccard dissimilarities (**Figure 1C**) (McInnes and Healy, 2018).

**Figure 1.**
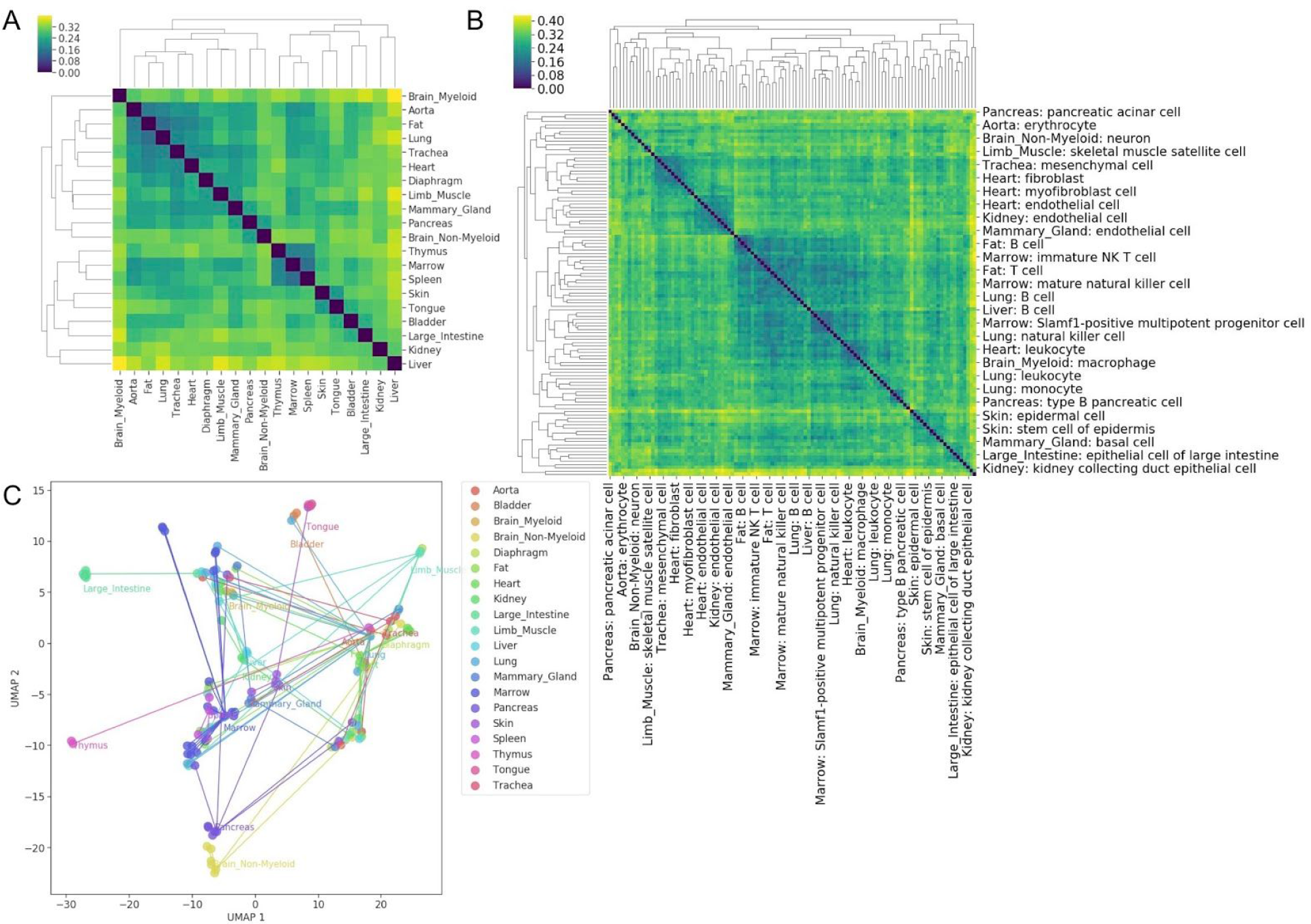
Tissue and cell type context-specific metabolic models constructed using Tabula Muris scRNA-seq dataset (Schaum et al., 2018). (**A**) Pairwise Jaccard dissimilarities between tissue type specific metabolic models. (**B**) Pairwise Jaccard dissimilarities between cell type specific metabolic models. (**C**) UMAP 2D embeddings of the tissue and cell type specific metabolic models using the Jaccard dissimilarities between their reaction indicators. The text labels are close to the tissue type specific models, and the lines connect tissue specific metabolic models and their corresponding cell type specific models.

The variations between different metabolic models showed that the metabolic modeling procedure is sensitive to the differences in transcriptomic profiles. The UMAP embeddings of metabolic models (**Figure 1C**) separated different cells in a similar pattern as the UMAP embeddings computed by the cosine distance of single-cell gene read counts of the metabolic genes in iMM1415 (**Figure S3B**). The UMAP embeddings of metabolic models (**Figure 2C**) and 1,375 metabolic genes (**Figure S3B**) are less capable of separating different tissue types, comparing to the UMAP embeddings of all 44,728 genes, which implies a less diverse space of cellular metabolic states compared to cell types. The mean number of reactions in all cell type models is 1,376, and the standard deviation is 76 (**Figure S4A**). The mean and standard deviation of the number of reactions in all tissue type models are 1,383 and 83 (**Figure S4A**).

**Figure 2.**
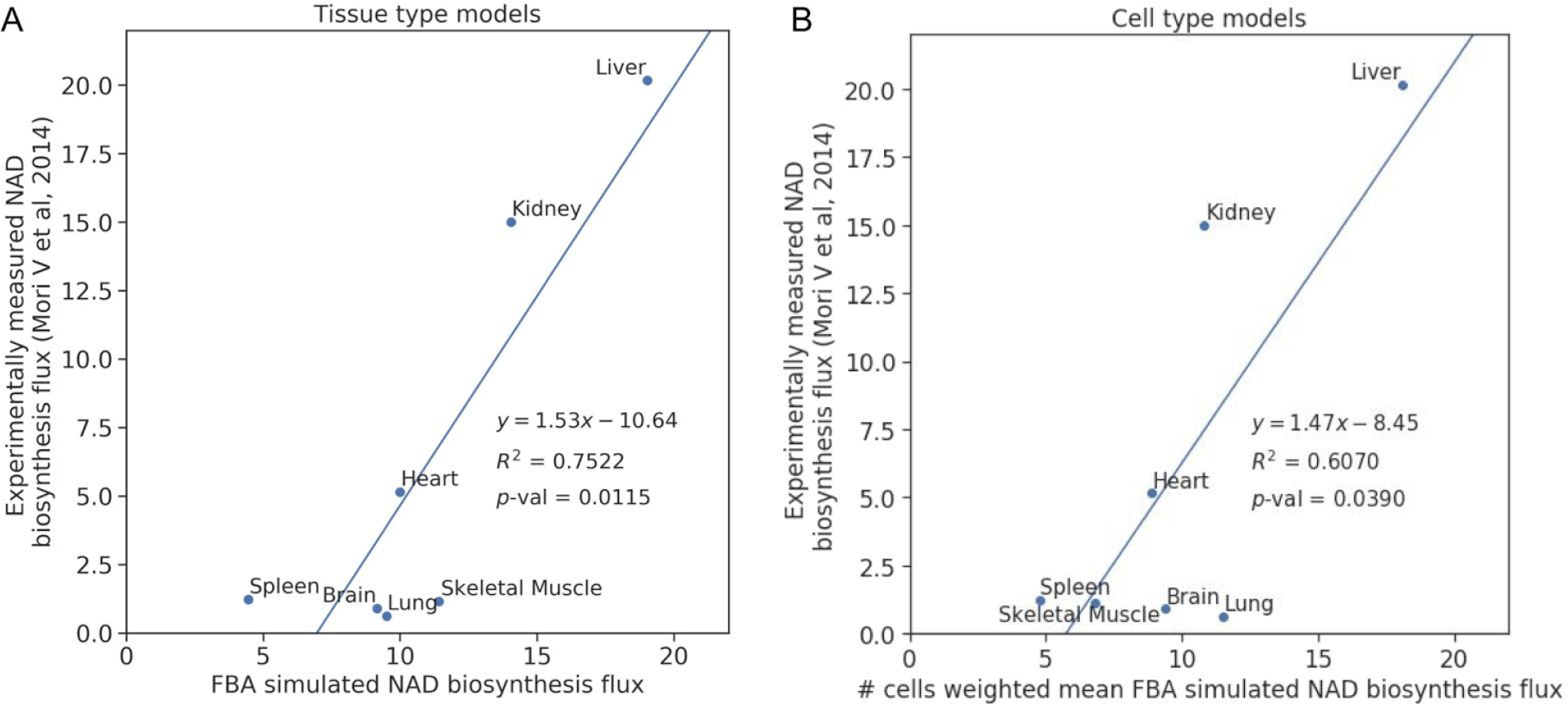
FBA simulated NAD^+^ biosynthesis fluxes. (**A**) Linear correlation between tissue type model simulated NAD^+^ biosynthesis fluxes and empirically measured ones. (**B**) Linear correlation between cell type model simulated NAD^+^ biosynthesis fluxes and empirically measured ones, in which the simulated tissue fluxes are the mean of cell type fluxes weighted by the number of cells in each tissue.

In order to deal with the high drop-out rate in scRNA-seq, we agglomerated multiple cells together to construct a metabolic model. The mean number of expressed genes in all cell types is 9,131, and the standard deviation is 1,490 (**Figure S4B**). In all tissue types, the mean number of expressed genes is 10,599, and the standard deviation is 993 (**Figure S4B**). The nearly 10,000 mean number of expressed genes suggests a low gene dropout rate with regard to cell types and tissue types. The median number of cells in all cell types and tissue types is 178 and 1,630 respectively (**Figure S4C**), which suggests that the probability is low for a gene to be stochastically dropped out in all cells of a cell or tissue type.

### Simulated tissue specific NAD^+^ biosynthesis fluxes

The simulated tissue specific NAD^+^ biosynthesis fluxes show significant (p < 0.05) linear correlation with empirically measured ones (Mori et al., 2014) (**Figure 2A** and **2B**). The R^2^ value is higher in tissue type model simulated fluxes, which may be caused by the inaccurate estimations of reaction confidence scores in certain cell types due to the low number of single cells in the dataset. When the number of cells is low, the estimation of gene expression confidence scores is strongly affected by the noise in scRNA-seq data. Also, the number of cells in each cell type of a tissue may not accurately represent the in vivo composition (Newman et al., 2019), which would result in the inaccurate weights in the computation of tissue specific fluxes using cell type model simulated fluxes.

The linear correlations between experimentally measured and FBA simulated NAD^+^ biosynthesis fluxes cannot be directly explained by the expression levels of individual genes. The linear correlations are not significant between empirically measured NAD^+^ biosynthesis fluxes and the expression levels of the enzymes that are catalyzing the last step of the NAD^+^ biosynthesis (**Figure S5**), including Nmnat1, Nmnat2, Nmnat3, and Nadsyn1 (**Table S1**). Although the expression levels of Qprt and Nmrk1 show significant linear correlations to the empirically measured NAD^+^ biosynthesis fluxes, the false discovery rates (FDRs) are not significant when Qprt and Nmrk1 are tested together with the key enzymes in NAD^+^ biosynthesis (**Table S1** and **Figure S5**) (Mori et al., 2014), all genes (**Table S3**), or metabolic model iMM1415 genes (**Table S4**). The lowest FDRs in the linear correlations between all 22,454 genes and empirically measured NAD^+^ biosynthesis fluxes is 0.071542. Although 72 genes in the iMM1415 metabolic model have FDR < 0.05, interpreting these significant linear correlations is not straightforward, and the key enzymes in NAD^+^ biosynthesis all have FDRs > 0.09.

### Metabolic variations within tissue and cell types estimated by bootstrapping

We estimated the metabolic variations within different tissue and cell types by bootstrapping. The standard deviations of bootstrapping sample fluxes are used to illustrate the extent of metabolic variations, and the means are used to compute the significance of linear correlation with empirically measured fluxes (**Figures 3A** and **3B**). The linear correlation significance varies in each bootstrapping run (**Figures S6B** and **S6C**), which may be caused by the signal-to-noise ratio in the bootstrapping samples. In addition, the bootstrapping fluxes provide the estimates of the metabolic variations (**Figures 3C** and **3D**), which is informative in exploring the heterogeneous scRNA-seq datasets.

**Figure 3.**
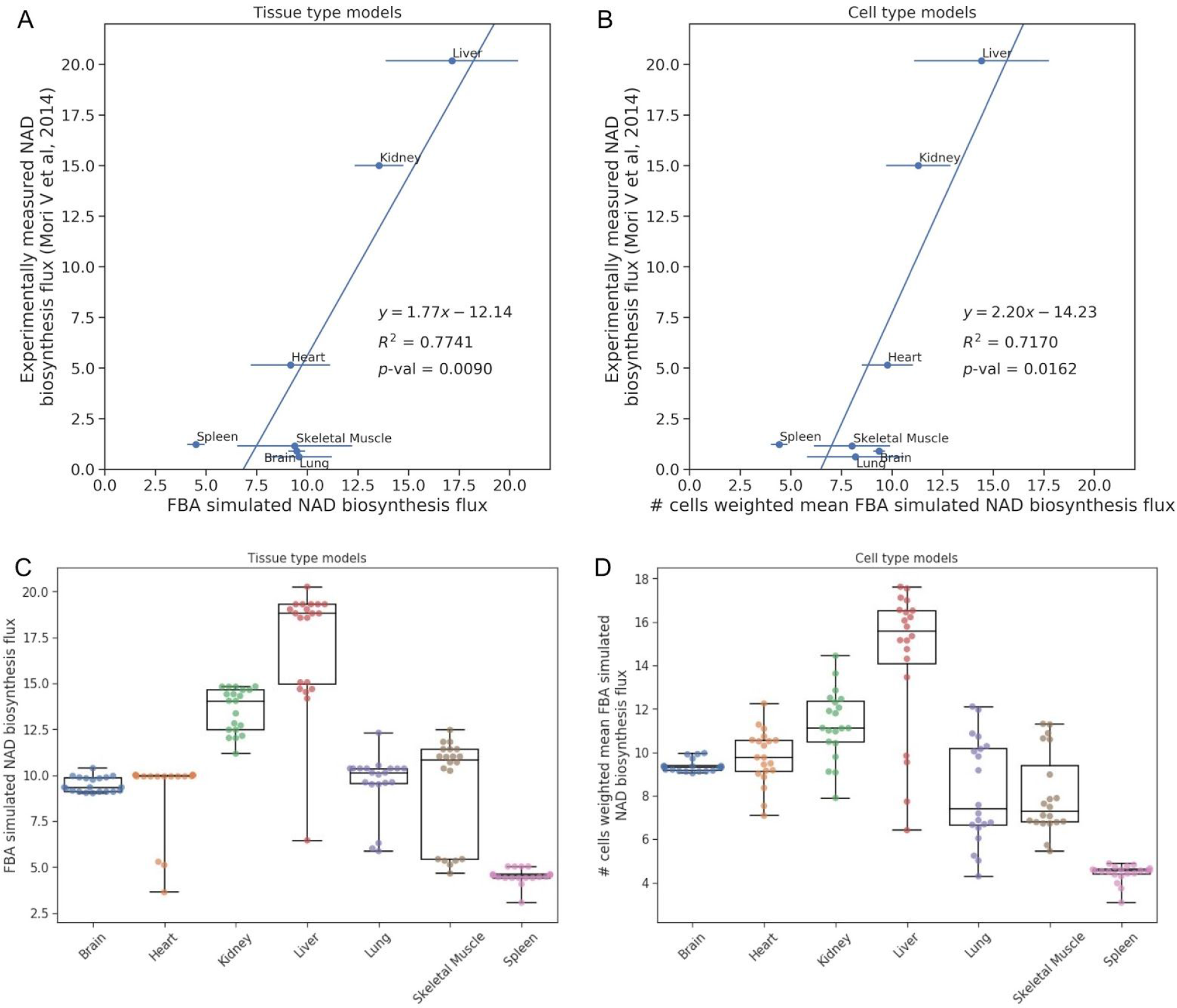
FBA simulated NAD^+^ biosynthesis fluxes with bootstrapping. (**A**) Linear correlation between tissue type model simulated NAD^+^ biosynthesis fluxes and empirically measured ones. The error bars are the standard deviation of the bootstrapping samples. (**B**) Linear correlation between cell type model simulated NAD^+^ biosynthesis fluxes and empirically measured ones, in which the simulated tissue fluxes are the mean of cell type fluxes weighted by the number of cells in each tissue. The error bars are the standard deviation of the bootstrapping samples. (**C**) Distribution of the tissue type model simulated NAD^+^ biosynthesis fluxes in all bootstrapping samples. (**D**) Distribution of the cell type model simulated NAD^+^ biosynthesis fluxes in all bootstrapping samples, in which the simulated tissue fluxes are the mean of cell type fluxes weighted by the number of cells in each tissue.

The distribution of NAD^+^ biosynthesis fluxes in bootstrapping samples implies that a tissue type may be composed of cells in multiple metabolic states (**Figure 3C** and **3D**). The fluxes are separated into high and low groups in several tissues, such as kidney and skeletal muscle. Similar separations are also shown in the fluxes of bootstrapping samples of several cell types (**Figure S6A**), such as kidney macrophage and lung endothelial cell.

### Simulated metabolic fluxes are reproducible using different scRNA-seq datasets

We constructed cell-type specific metabolic models using the scRNA-seq datasets from Allen Brain Mouse Atlas (ABMA), and we simulated the NAD^+^ biosynthesis fluxes using the constructed cell-type specific models (**Figure 4A**). We found the simulated NAD^+^ biosynthesis fluxes using Tabula Muris scRNA-seq dataset are reproducible using ABMA ALM and VISp scRNA-seq datasets (**Figure 4A** and **Table S2**). The simulated NAD^+^ biosynthesis fluxes are similar among the common cell types in the three different scRNA-seq datasets. Although the simulated NAD^+^ biosynthesis fluxes tend to have the same ascending order of ABMA ALM < ABMA VISp < Tabula Muris in the same cell type, the order of the simulated NAD^+^ biosynthesis fluxes of different cell types within a dataset is relatively consistent across the three scRNA-seq datasets.

**Figure 4.**
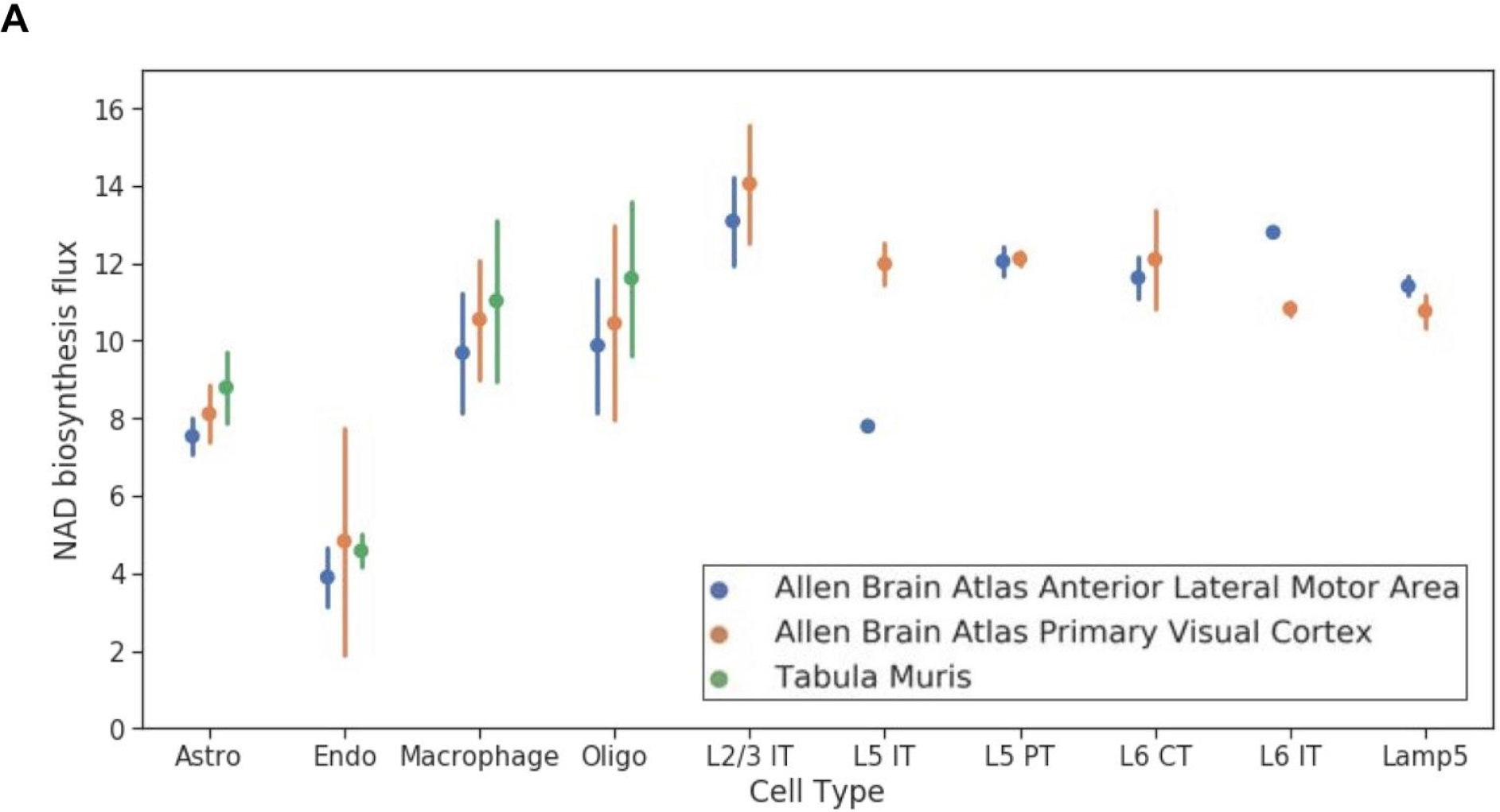
Simulated cell-type specific NAD^+^ biosynthesis fluxes using scRNA-seq datasets from Tabula Muris and Allen Brain Mouse Atlas. (**A**) Reproducibility of metabolic model NAD^+^ biosynthesis flux from iMM1415 using CORDA. Results are shown for mean NAD^+^ biosynthesis flux by selected cell types from Tabula Muris, Allen Brain Institute (ABI): Anterior Lateral Motor Area (ALM) and ABI Primary Visual Cortex (VISp) scRNAseq. The points and error bars represent the mean and standard deviation (sd) flux values for NAD^+^ biosynthesis simulated by 20 bootstrapping replicates at 80% of the cells per set. Cell types include astrocytes (Astro), endothelial cells (Endo), oligodendrocytes (Oligo) and neurons indicated by cortical layers 2-5 (L2-L5; L4 is not included in bootstrapping with only 3 cells). Intratelencephalic (IT), pyramidal tract (PT), and corticothalamic (CT). Lamp5: subclass of GABAergic neurons.

## Discussion

The biological interpretations of scRNA-seq results are obtained from extensive data analyses, which could take more time than doing the experiments. As the size of scRNA-seq datasets increases rapidly (Rozenblatt-Rosen et al., 2017; Svensson et al., 2018), the scRNA-seq profiled biological processes become increasingly complex, which involve multiple cell types, tissue types, and organs that are undergoing rapid morphological and molecular changes (Cao et al., 2019; Haldipur et al., 2019; Schiebinger et al., 2019). In such complex biological processes, interpreting the single-cell transcriptomic changes requires integration of relevant regulatory mechanisms of gene expression and cell state transition (Tanay and Regev, 2017).

The results of scRNA-seq provide rough measurements of expressed genes in each cell, but the measurements need to be interpreted with extra biological understanding. As biological processes generally involve multiple interacting components, a comprehensive interpretation of single-cell transcriptomic profiles requires a systematic framework that integrates all relevant biological components and their interactions. The systematic frameworks can be developed by mathematically modeling the biological components and their interactions that are comprehensively curated and structured in databases. The previously developed frameworks for interpreting RNA microarray and bulk RNA-seq datasets cannot be directly adapted to interpret scRNA-seq results because of the unique statistical properties of scRNA-seq data and biological purposes of scRNA-seq experiments. Generally, scRNA-seq data have lower signal-to-noise ratio, and scRNA-seq experiments aim to uncover the cellular heterogeneity and identify rare cell populations within cell and tissue types.

We developed a computational method to systematically interpret scRNA-seq dataset in the aspect of cellular metabolism. The method integrates the databases of metabolic reactions (Brunk et al., 2018; Kanehisa et al., 2016; King et al., 2016) to infer metabolic flux variations between different cell and tissue types using scRNA-seq data, through constraint based metabolic modeling (Ebrahim et al., 2013), which is more reliable than associating flux variations with individual gene expression levels (Hackett et al., 2016). In a constraint based metabolic model, the metabolites and reactions are represented as either discrete mathematical graphs (**Figure S1A)** or a system of linear equations (**Figure S1B**). In the graph representation, each metabolite is a node, and each reaction is an edge, which can be intuitively used for visualization. In the linear system representation, each reaction is an equation with reactants on the left hand side and products on the right, which can be used for simulating metabolic fluxes using optimization and sampling approaches, such as FBA (**Figure S2**) and uniform sampling (Megchelenbrink et al., 2014). The reference model containing all metabolic reactions of an organism can be reduced to represent cell- and tissue-type specific metabolism, according to the scRNA-seq profiled transcription levels of enzymes that are catalyzing the reactions. In order to handle the relatively low signal-to-noise ratio in scRNA-seq data, we agglomerated cells with the same cell or tissue type together to obtain a more accurate transcriptomic profile, and the cellular heterogeneity within the cell and tissue types are evaluated by bootstrapping. The transcriptomic levels are also discretized into five levels to increase the robustness of the modeling and simulation methods.

Computational modeling of cell- and tissue-type specific metabolism using scRNA-seq data is able to guide the design of studies to uncover the regulatory mechanisms of cellular metabolism. Experimental measurements of cellular metabolism are mainly performed by specialized biochemical assays, mass spectrometry, and nuclear magnetic resonance spectrometry (Dunn et al., 2011). As such experiments take a considerable amount of time and resources (Jang et al., 2018), mathematical modeling of the metabolic fluxes would be able to guide a careful experiment design to facilitate the progression of a study.

The developed metabolic modeling method can further be applied to study epigenetic reprogramming and metabolite deficiency. Biosynthetic fluxes of substrates used for epigenetic modifications can be simulated to infer whether cells are undergoing active epigenetic reprogramming in various differentiation or development processes (Reid et al., 2017). Biosynthetic fluxes of metabolites involved in cell survival can be simulated to determine whether the cells have sufficient bioenergetics in their supply of metabolites (Williams et al., 2017).

The constraint based mathematical modeling strategy of metabolism can be further applied to model other biological processes, through representing relevant biological components and events as networks and linear equation systems. Depending on the complexity of the modeled biological process, different levels of abstraction needs to be applied to ensure the feasibility of modeling procedure. In this way, multiple modeling frameworks can further be integrated to interpret multi-omics datasets.

## References

Bordbar, A., Monk, J.M., King, Z.A., and Palsson, B.O. (2014). Constraint-based models predict metabolic and associated cellular functions. Nat. Rev. Genet. 15, 107–120.

Brunk, E., Sahoo, S., Zielinski, D.C., Altunkaya, A., Dräger, A., Mih, N., Gatto, F., Nilsson, A., Preciat Gonzalez, G.A., Aurich, M.K., et al. (2018). Recon3D enables a three-dimensional view of gene variation in human metabolism. Nat. Biotechnol. 36, 272–281.

Cao, J., Spielmann, M., Qiu, X., Huang, X., Ibrahim, D.M., Hill, A.J., Zhang, F., Mundlos, S., Christiansen, L., Steemers, F.J., et al. (2019). The single-cell transcriptional landscape of mammalian organogenesis. Nature 1.

Childs, B., Valle, D., and Jimenez-Sanchez, G. (2001). The inborn error and biochemical individuality. The Metabolic and Molecular Bases of Inherited Disease 1, 155–166.

DeBerardinis, R.J., and Thompson, C.B. (2012). Cellular metabolism and disease: what do metabolic outliers teach us? Cell 148, 1132–1144.

Dunn, W.B., Broadhurst, D.I., Atherton, H.J., Goodacre, R., and Griffin, J.L. (2011). Systems level studies of mammalian metabolomes: the roles of mass spectrometry and nuclear magnetic resonance spectroscopy. Chem. Soc. Rev. 40, 387–426.

Ebrahim, A., Lerman, J.A., Palsson, B.O., and Hyduke, D.R. (2013). COBRApy: COnstraints-Based Reconstruction and Analysis for Python. BMC Syst. Biol. 7, 74.

Hackett, S.R., Zanotelli, V.R.T., Xu, W., Goya, J., Park, J.O., Perlman, D.H., Gibney, P.A., Botstein, D., Storey, J.D., and Rabinowitz, J.D. (2016). Systems-level analysis of mechanisms regulating yeast metabolic flux. Science 354.

Haldipur, P., Aldinger, K.A., Bernardo, S., Deng, M., Timms, A.E., Overman, L.M., Winter, C., Lisgo, S.N., Razavi, F., Silvestri, E., et al. (2019). Spatiotemporal expansion of primary progenitor zones in the developing human cerebellum. Science.

Jang, C., Chen, L., and Rabinowitz, J.D. (2018). Metabolomics and Isotope Tracing. Cell 173, 822–837.

Kanehisa, M., Sato, Y., Kawashima, M., Furumichi, M., and Tanabe, M. (2016). KEGG as a reference resource for gene and protein annotation. Nucleic Acids Res. 44, D457–D462.

King, Z.A., Lu, J., Dräger, A., Miller, P., Federowicz, S., Lerman, J.A., Ebrahim, A., Palsson, B.O., and Lewis, N.E. (2016). BiGG Models: A platform for integrating, standardizing and sharing genome-scale models. Nucleic Acids Res. 44, D515–D522.

Leone, R.D., Zhao, L., Englert, J.M., Sun, I.-M., Oh, M.-H., Sun, I.-H., Arwood, M.L., Bettencourt, I.A., Patel, C.H., Wen, J., et al. (2019). Glutamine blockade induces divergent metabolic programs to overcome tumor immune evasion. Science 366, 1013–1021.

Mahadevan, R., and Schilling, C.H. (2003). The effects of alternate optimal solutions in constraint-based genome-scale metabolic models. Metab. Eng. 5, 264–276.

McInnes, L., and Healy, J. (2018). UMAP: Uniform Manifold Approximation and Projection for Dimension Reduction.

Megchelenbrink, W., Huynen, M., and Marchiori, E. (2014). optGpSampler: an improved tool for uniformly sampling the solution-space of genome-scale metabolic networks. PLoS One 9, e86587.

Mori, V., Amici, A., Mazzola, F., Di Stefano, M., Conforti, L., Magni, G., Ruggieri, S., Raffaelli, N., and Orsomando, G. (2014). Metabolic profiling of alternative NAD biosynthetic routes in mouse tissues. PLoS One 9, e113939.

Newman, A.M., Steen, C.B., Liu, C.L., Gentles, A.J., Chaudhuri, A.A., Scherer, F., Khodadoust, M.S., Esfahani, M.S., Luca, B.A., Steiner, D., et al. (2019). Determining cell type abundance and expression from bulk tissues with digital cytometry. Nat. Biotechnol.

Orth, J.D., Thiele, I., and Palsson, B.Ø. (2010). What is flux balance analysis? Nat. Biotechnol. 28, 245–248.

Pacheco, M.P., Pfau, T., and Sauter, T. (2015). Benchmarking Procedures for High-Throughput Context Specific Reconstruction Algorithms. Front. Physiol. 6, 410.

Pierre, G. (2013). Neurodegenerative disorders and metabolic disease. Arch. Dis. Child. 98, 618–624.

Potter, M., Newport, E., and Morten, K.J. (2016). The Warburg effect: 80 years on. Biochem. Soc. Trans. 44, 1499–1505.

Reid, M.A., Dai, Z., and Locasale, J.W. (2017). The impact of cellular metabolism on chromatin dynamics and epigenetics. Nat. Cell Biol. 19, 1298–1306.

Rozenblatt-Rosen, O., Stubbington, M.J.T., Regev, A., and Teichmann, S.A. (2017). The Human Cell Atlas: from vision to reality. Nature 550, 451–453.

Schaum, N., Karkanias, J., Neff, N.F., May, A.P., Quake, S.R., Wyss-Coray, T., Darmanis, S., Batson, J., Botvinnik, O., Chen, M.B., et al. (2018). Single-cell transcriptomics of 20 mouse organs creates a Tabula Muris. Nature.

Schellenberger, J., and Palsson, B.Ø. (2009). Use of randomized sampling for analysis of metabolic networks. J. Biol. Chem. 284, 5457–5461.

Schiebinger, G., Shu, J., Tabaka, M., Cleary, B., Subramanian, V., Solomon, A., Gould, J., Liu, S., Lin, S., Berube, P., et al. (2019). Optimal-Transport Analysis of Single-Cell Gene Expression Identifies Developmental Trajectories in Reprogramming. Cell 176, 1517.

Schultz, A., and Qutub, A.A. (2016). Reconstruction of Tissue-Specific Metabolic Networks Using CORDA. PLoS Comput. Biol. 12, e1004808.

Sigurdsson, M.I., Jamshidi, N., Steingrimsson, E., Thiele, I., and Palsson, B.Ø. (2010). A detailed genome-wide reconstruction of mouse metabolism based on human Recon 1. BMC Syst. Biol. 4, 140.

Svensson, V., Vento-Tormo, R., and Teichmann, S.A. (2018). Exponential scaling of single-cell RNA-seq in the past decade. Nat. Protoc. 13, 599–604.

Tanay, A., and Regev, A. (2017). Scaling single-cell genomics from phenomenology to mechanism. Nature 541, 331–338.

Warburg, O., Wind, F., and Negelein, E. (1927). THE METABOLISM OF TUMORS IN THE BODY. J. Gen. Physiol. 8, 519–530.

Williams, P.A., Harder, J.M., Foxworth, N.E., Cochran, K.E., Philip, V.M., Porciatti, V., Smithies, O., and John, S.W.M. (2017). Vitamin B3 modulates mitochondrial vulnerability and prevents glaucoma in aged mice. Science 355, 756–760.

