## Supplementary Methods, Figures and Tables for "Modeling metabolic variation with single-cell expression data"

##### *Context-specific metabolic model construction using scRNA-seq data*

Modeling cellular metabolism is motivated by the identification of various metabolic pathways in cells such as glycolysis, tricarboxylic acid (TCA) cycle, and oxidative phosphorylation. In the Recon3D curated model of human metabolism, there are 5,835 metabolites and 10,600 reactions (Brunk et al., 2018). In addition to the large number of metabolites and reactions, the reactions interact in a non-independent and non-additive manner (Le Novère, 2015), and the metabolites are extensively consumed in other biological processes, such as nucleotides in transcription, amino acids in translation, $NAD^+$ in parylation (Gibson and Kraus, 2012), and acetyl-CoA in histone acetylation (Reid et al., 2017). Furthermore, metabolites are constantly exchanged across cellular membranes through transporters and diffusion. Mathematically describing the current knowledge of cellular metabolism is able to facilitate experiment design and interpretation of results.

Mathematical modeling of cellular metabolism is based on the principles of kinetics in physical chemistry. The relationships between metabolite concentrations and reaction fluxes are written as a system of differential equations as the following (Costa et al., 2016),

$$\frac{d[X_i]}{dt} = \sum_j \mathbf{S}_{i,j} \cdot v_j([E], [X], \vec{k}).$$

In the differential equations:

- $[X_i]$ is the concentration of the $i$-th metabolite
- $t$ is time.
- $\mathbf{S}$ is the stoichiometric matrix of the metabolic model (**Figure S1C**). Each row represents a metabolite, and each column represents a reaction. The entry $\mathbf{S}_{i,j}$ is the stoichiometric coefficient of the $i$-th metabolite in $j$-th reaction. $\mathbf{S}_{i,j}$ is positive if the $i$-th metabolite is on the right-hand side of the reaction, otherwise negative. For example, the 3rd reaction in **Figure S1B** is "A[cyt] + B[cyt] → C[cyt]", so $\mathbf{S}_{1,3} = -1$, $\mathbf{S}_{4,3} = -1$, and $\mathbf{S}_{5,3} = 1$.

- $v_j$ is the reaction flux, or rate, of the $j$-th reaction, converting left-hand-side reactants into right-hand-side products. Fluxes are dependent on the concentrations of metabolites ($[X]$) and enzymes ($[E]$) and the kinetic constants of the enzymes ($\vec{k}$). The enzyme concentrations can be considered as constants within a certain period of time or variables dependent on parameters relevant to protein degradation. When the reaction has only one reactant and one product and is catalyzed by one enzyme, the flux of the reaction can be computed with
$$v = [E] \cdot k_{cat} \cdot \frac{[S]}{[S] + K_m},$$
where $[E]$ and $[S]$ are the concentrations of the enzyme and substrate respectively. $k_{cat}$ is turnover number, which is a kinetic constant of the enzyme representing the number of catalytic cycles can be performed by the enzyme within a unit of time. $K_m$ is Michaelis constant, which is a constant representing the affinity between the enzyme and its substrate.

The differential equation system defines the changes of metabolite concentrations and reaction fluxes over time, given initial concentrations of metabolites and enzymes, stoichiometric coefficients, and kinetic parameters (Saa and Nielsen, 2017).

There are mainly two methods to use the differential equation system to model cellular metabolism, either kinetic modeling or constraint-based modeling. In kinetic modeling, experimentally measured $[X]$ and $\vec{k}$ of a cell type are plugged into the differential equation system, and the changes of $[X]$ and $v$ are simulated over time to characterize the metabolism of the cell type. In constraint-based modeling, the differential equation system is analyzed under steady metabolite concentration state, in which the changes of metabolite concentrations are all equal to 0. The steady metabolite concentration state is biologically meaningful, because the changes of metabolite concentrations are relatively slower than the changes of metabolic fluxes (Edwards and Palsson, 2000; Fell, 2005; Palsson and Lightfoot, 1984). Under steady state, the differential equation system becomes a system of linear equations with the fluxes as variables, as the following
$$\sum_j \mathbf{S}_{i,j} \cdot v_j = 0.$$
Although $v_j$ can still be computed by kinetic laws given $([E], [X], \vec{k})$, we can compute $v_j$ without $([E], [X], \vec{k})$ by assuming that cellular metabolism optimizes the fluxes of a set of reactions, such as the reactions involved in cell growth, through linear optimization with additional lower and upper bounds of reaction fluxes. The lower and upper bounds of reaction fluxes are called constraints in a linear programming problem, and the bounds can be experimentally determined or computationally estimated (Hackett et al., 2016).

We selected constraint-based modeling to infer cell-type specific metabolic fluxes using scRNA-seq data, because constraint-based modeling requires less initial information than kinetic modeling. Kinetic modeling requires kinetic constants of enzymes and concentrations of metabolites and enzymes that are not available for mouse or human, whereas constraint-based modeling only requires the stoichiometric coefficients and flux constraints. Although the flux constraints are also not available for different mouse or human cell types, the computed fluxes through constraint-based modeling are robust to inaccurate flux constraints due to the robustness and redundancy of metabolic networks (Edwards and Palsson, 2000).

In constraint-based metabolic flux computation, the cell-type specificity is determined by the enzyme transcription levels in different cell types characterized by scRNA-seq. We infer whether an enzyme is expressed in a cell type or not from scRNA-seq data, based on the average read counts of the enzyme in the single cells with the same cell type. Considering that enzymes are transcribed in vivo at distinct levels, we applied CORDA to refine the determination of whether an enzyme is expressed or not in a cell type (Schultz and Qutub, 2016), through a linear programming setting to minimize the inclusion of lowly transcribed enzymes while favoring the inclusion of important enzymes. The enzymes that are determined to be expressed in the cell type formulate a cell-type specific stoichiometric matrix $\mathbf{S}$, which is further used for flux computation in various optimization settings. The upper and lower bounds of the fluxes are not customized for the cell type based on the enzyme transcription levels, but such procedures can be integrated in the future to improve the sensitivity of flux computation to distinct transcriptomic profiles.

The constraint-based flux computation is either based on optimization or sampling procedures under steady state. In optimization procedures, flux balance analysis (FBA) is the foundation of various other methods. In FBA, we assume that cellular metabolism under steady state is optimized for certain objectives, such as cell growth and ATP production. Therefore, the metabolic differential equation system can be written as a linear programming problem (**Figure S2**), and the problem can be efficiently solved in polynomial time using free or commercial software to obtain the maximum values of objective fluxes (Meindl and Templ, 2012). The maximum objective fluxes are cell-type specific, because different cell types have different stoichiometric matrix $\mathbf{S}$. In sampling procedures, metabolic fluxes are sampled from the allowable solution space defined by steady state $\mathbf{S} \cdot \mathbf{v} = \mathbf{0}$ and flux constraints $\mathbf{l} \leq \mathbf{v} \leq \mathbf{u}$ based on certain statistical distributions (Megchelenbrink et al., 2014).

#### *Exchange, demand and sink reactions*

Constraint based metabolic models are open chemical systems, which are able to import metabolites from external chemical systems and export metabolites to external chemical systems. The interactions between the models and the external chemical systems are mediated through exchange, demand, and sink reactions. Such reactions are written as mass unbalanced reactions in the models (**Figure S1B**), but they are actually balanced with the external chemical systems. The exchange, demand, and sink reactions are also biologically meaningful.

The exchange reactions are reversible reactions with the metabolites in extracellular space as reactants, which represent the availability and consumability of metabolites in the external chemical system. The exchange reactions are defined by the culturing medium or extracellular environment of the modeled cells. The extracellular environment is distinct from the extracellular space in constraint based models (**Figure S1A**). The metabolites and reactions in extracellular space are internal to the model, representing the interactions between the modeled cells and their extracellular space. The extracellular environment is external to the model. The exchange reactions transport metabolites between extracellular space and extracellular environment, which define the availability of metabolites for the modeled cell (Sigurdsson et al., 2010; Thiele and Palsson, 2010). If an exchange reaction carries a positive flux, it means its corresponding reactant is exported from the extracellular space into the extracellular environment. If an exchange reaction carries a negative flux, it means its corresponding reactant is imported from the extracellular environment to the extracellular space.

The demand reactions are irreversible reactions with the metabolites inside the modeled cells as reactants (Thiele and Palsson, 2010), which represent the accumulation or consumption of metabolites inside the modeled cells. Conventionally, the direction of an irreversible reaction is from the left hand side of the formula to the right hand side, so irreversible reactions can only carry non-negative fluxes. If a demand reaction carries a positive flux, it means the reactant is accumulated in the external chemical system or consumed by the external chemical system. We optimize demand reactions to simulate the biosynthesis of metabolites.

The sink reactions are reversible reactions with the metabolites inside the modeled cells as reactants (Thiele and Palsson, 2010), which have different meanings when carrying positive or negative fluxes. If a sink reaction carries positive fluxes, it has the same meaning as a demand reaction with the same flux. If a sink reaction carries negative flux, it means that the reactant is provided by the external chemical system.

The definitions of exchange, demand and sink reactions may be slightly different in different publications, but their usage is the same, which is to enable interactions between the constraint based metabolic models and their external chemical systems.

##### *Jaccard dissimilarity between two context specific metabolic models*

For each context specific metabolic model, we construct an indicator vector to represent the presence and absence of each reaction in the reference metabolic model. If the i-th reaction is present in the context specific model, the i-th entry of the indicator vector has value 1. If the i-th reaction is absent in the context specific model, the i-th entry of the indicator vector has value 0. The Jaccard dissimilarity between two context specific metabolic models is computed as the Jaccard dissimilarity of their reaction indicator vectors, as the following

$$\frac{c_{10} + c_{01}}{c_{11} + c_{01} + c_{10}},$$

where $c_{ab}$ is the count of occurrences of the i-th entry of the first reaction indicator vector is equal to $a$ and the i-th entry of the second reaction indicator vector is equal to $b$ for i ≤ the number of total reactions (Kosub, 2019).

#### Supplementary figures

Example metabolic model

A. Network representation

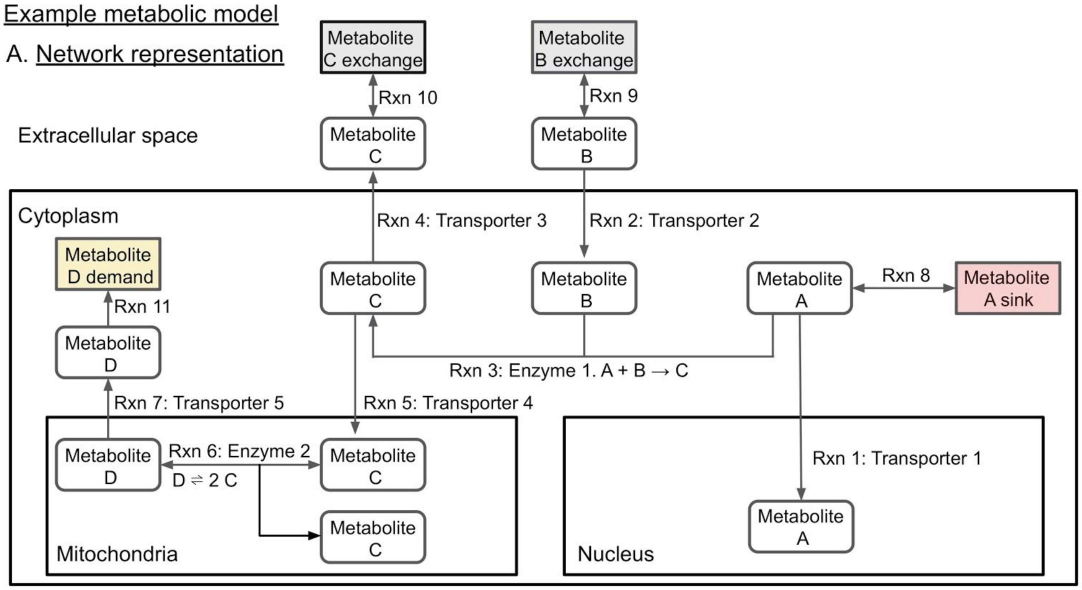


B. Reactions

| Reaction # and formula | Gene |
|---|---|
| 1: A[cyt] → A[nuc] | Transporter 1 |
| 2: B[ex] → B[cyt] | Transporter 2 |
| 3: A[cyt] + B[cyt] → C[cyt] | Enzyme 1 |
| 4: C[cyt] → C[ex] | Transporter 3 |
| 5: C[cyt] → C[mito] | Transporter 4 |
| 6: D[mito] ⇌ 2 C[mito] | Enzyme 2 |
| 7: D[mito] → D[cyt] | Transporter 5 |
| 8: A[cyt] ⇌ | N/A |
| 9: B[ex] ⇌ | N/A |
| 10: C[ex] ⇌ | N/A |
| 11: D[cyt] → | N/A |

C. Stoichiometric matrix

*Reactions* (columns) / *Metabolites* (rows)

| Metabolites | 1 | 2 | 3 | 4 | 5 | 6 | 7 | 8 | 9 | 10 | 11 |
|---|---|---|---|---|---|---|---|---|---|---|---|
| 1:A[cyt] | −1 | 0 | −1 | 0 | 0 | 0 | 0 | −1 | 0 | 0 | 0 |
| 2:A[nuc] | 1 | 0 | 0 | 0 | 0 | 0 | 0 | 0 | 0 | 0 | 0 |
| 3:B[ex] | 0 | −1 | 0 | 0 | 0 | 0 | 0 | 0 | −1 | 0 | 0 |
| 4:B[cyt] | 0 | 1 | −1 | 0 | 0 | 0 | 0 | 0 | 0 | 0 | 0 |
| 5:C[cyt] | 0 | 0 | 1 | −1 | −1 | 0 | 0 | 0 | 0 | 0 | 0 |
| 6:C[ex] | 0 | 0 | 0 | 1 | 0 | 0 | 0 | 0 | 0 | −1 | 0 |
| 7:C[mito] | 0 | 0 | 0 | 0 | 1 | 2 | 0 | 0 | 0 | 0 | 0 |
| 8:D[mito] | 0 | 0 | 0 | 0 | 0 | −1 | −1 | 0 | 0 | 0 | 0 |
| 9:D[cyt] | 0 | 0 | 0 | 0 | 0 | 0 | 1 | 0 | 0 | 0 | −1 |

**Figure S1.** Example metabolic model. (**A**) Network representation of the example metabolic model. There are four compartments in this example, including extracellular space, cytoplasm, mitochondria, and nucleus. Metabolites are transported from one compartment to another by transporters. Enzymes catalyze the conversions between metabolites. Reactions can be either irreversible or reversible. Three types of reactions are mass unbalanced, including exchange, demand and sink reactions. The metabolite demand is capable of consuming cytosolic metabolite D, and the metabolite D demand reaction export cytosolic metabolite D outside of the model. The metabolite A sink is capable of providing or consuming cytosolic metabolite A, and the metabolite A sink reaction imports cytosolic metabolite A into the model or exports cytosolic metabolite A outside of the model. The metabolite exchange reactions are capable of importing metabolites into the extracellular space from the external environment or exporting metabolites in the extracellular space into the external environment. (see Method section for more details about exchange, demand and sink reactions) (**B**) Reactions in the metabolic model. The text in the brackets represents the compartment of the metabolite: [ex] means extracellular space; [cyt] means cytoplasm; [nuc] means nucleus; [mito] meas mitochondria. The color shading represents the correspondence between the reaction formulae and stoichiometric coefficients of the metabolites. (**C**) Stoichiometric matrix $\mathrm{S}$ of the metabolic model. Each entry represents the stoichiometric coefficient of its corresponding metabolite (row) in its corresponding reaction (column) (see Supplementary Methods section for details).

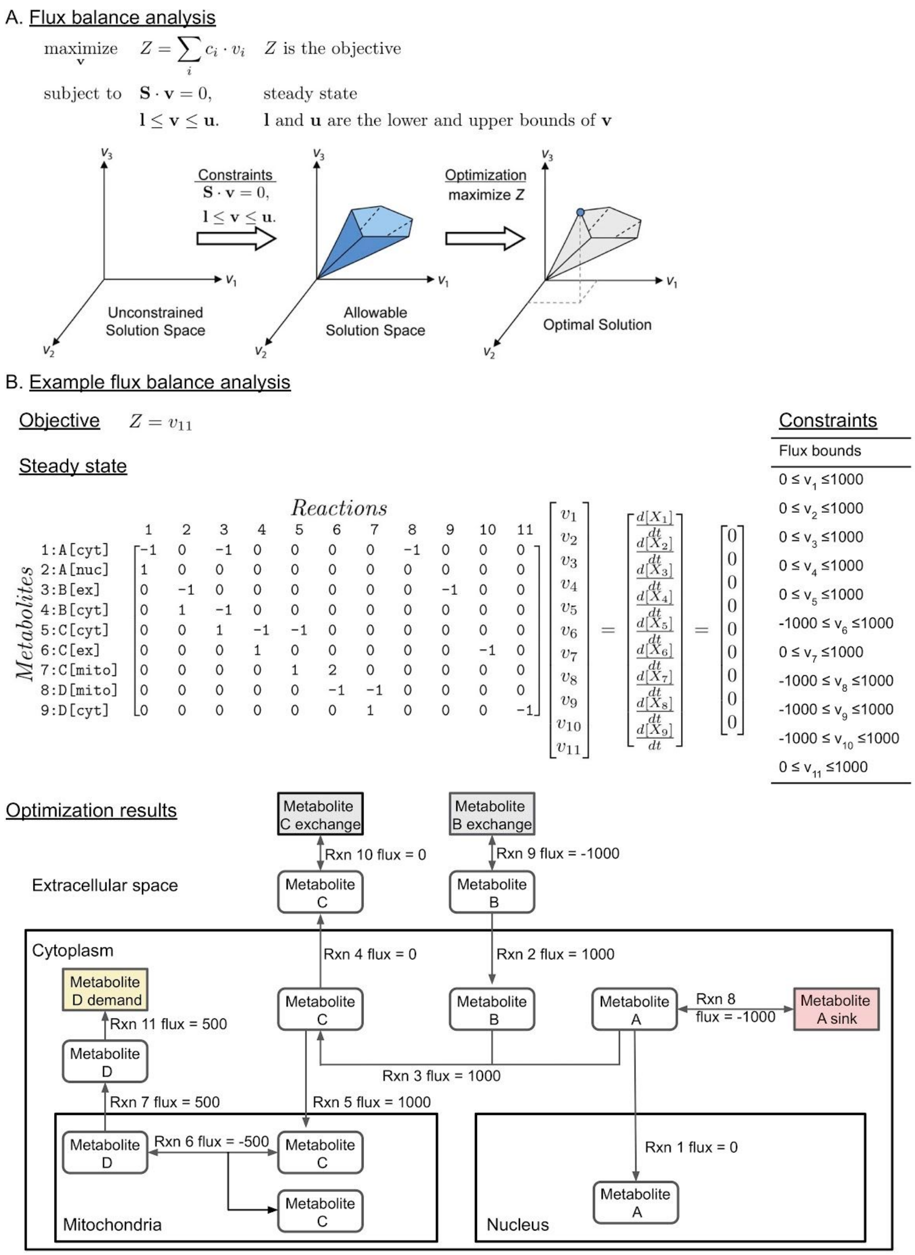


**Figure S2**. Flux balance analysis. (**A**) Flux balance analysis procedure, adapted from the illustration by Orth et al. (Orth et al., 2010). $Z$ is the objective of the linear programming problem, and it is a linear combination of the fluxes of one or more metabolic reactions. $\mathbf{S} \cdot \mathbf{v} = \mathbf{0}$ describes steady state, in which the metabolite concentrations do not change over time. $\mathbf{l} \leq \mathbf{v} \leq \mathbf{u}$ describes the flux constraints, so that the optimized flux values will not be infinite. (**B**) Example flux balance analysis on the metabolic model in **Figure S1A**. The objective is to maximize the flux of reaction 11, which is the demand reaction of metabolite D. The steady state is described as the equation $\mathbf{S} \cdot \mathbf{v} = \mathbf{0}$, which can be converted to a system of linear equations. The constraints are the reaction flux bounds, which are assigned according to the directionality of reactions. The optimization results are shown on the network representation of the metabolic model.

A

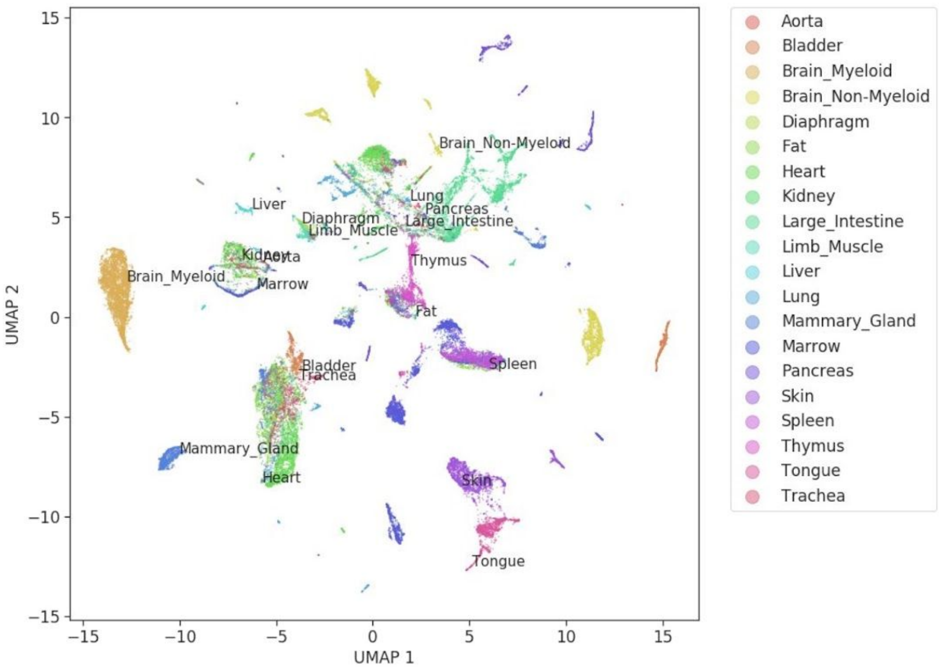


B

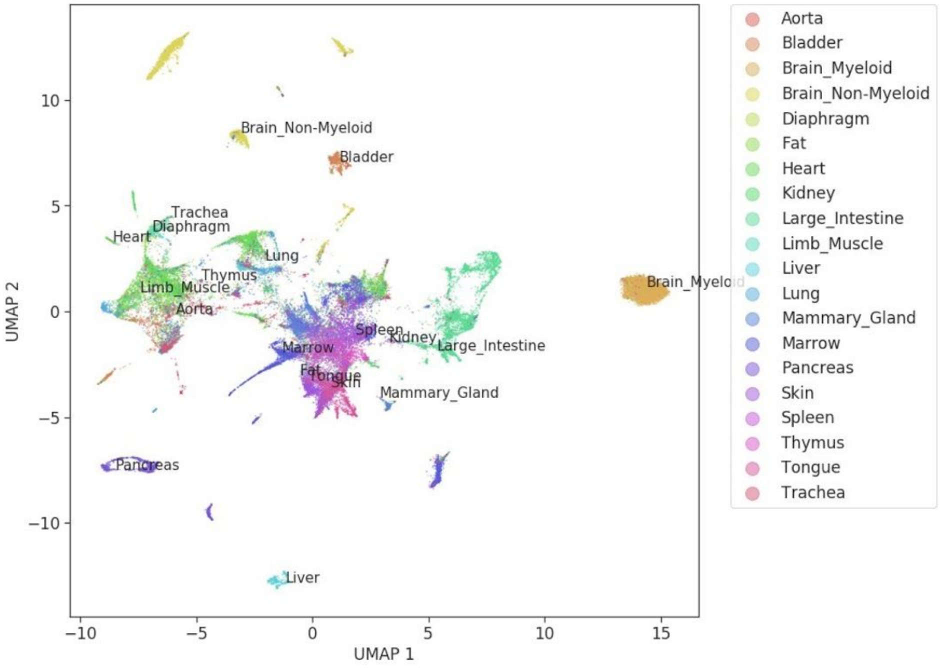


**Figure S3**. UMAP 2D embeddings of the transcriptomic profiles of different cells in Tabula Muris dataset (Schaum et al., 2018). (**A**) UMAP embeddings computed using all genes. (**B**) UMAP embeddings computed using only genes in the iMM1415 metabolic model.

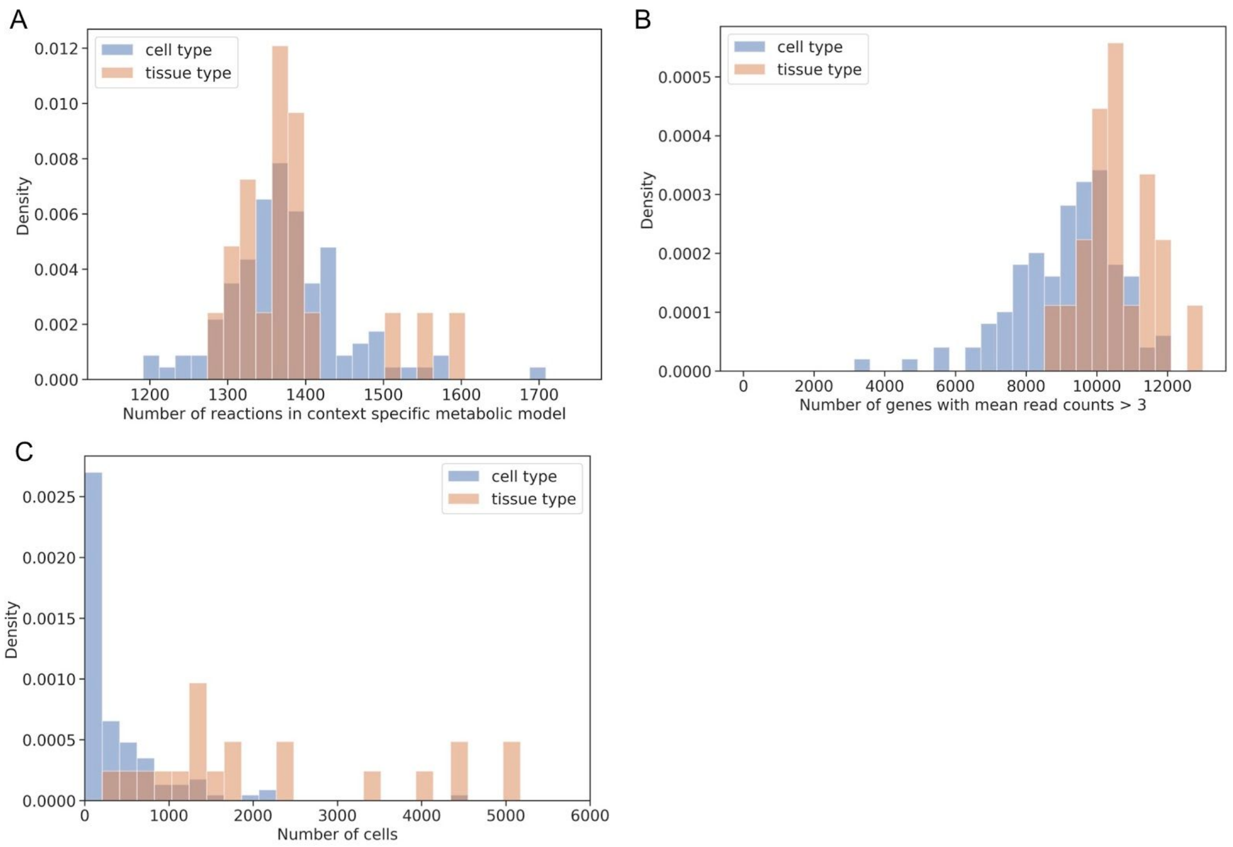


**Figure S4.** General statistics of the Tabula Muris scRNA-seq dataset and metabolic models. (**A**) Histogram of the number of reactions in cell- and tissue- type specific metabolic models constructed using the Tabula Muris dataset. (**B**) Histogram of the number of genes with mean read counts > 3 in different cell and tissue types in Tabula Muris dataset. (**C**) Histogram of the number of cells in different cell and tissue types in Tabula Muris dataset.

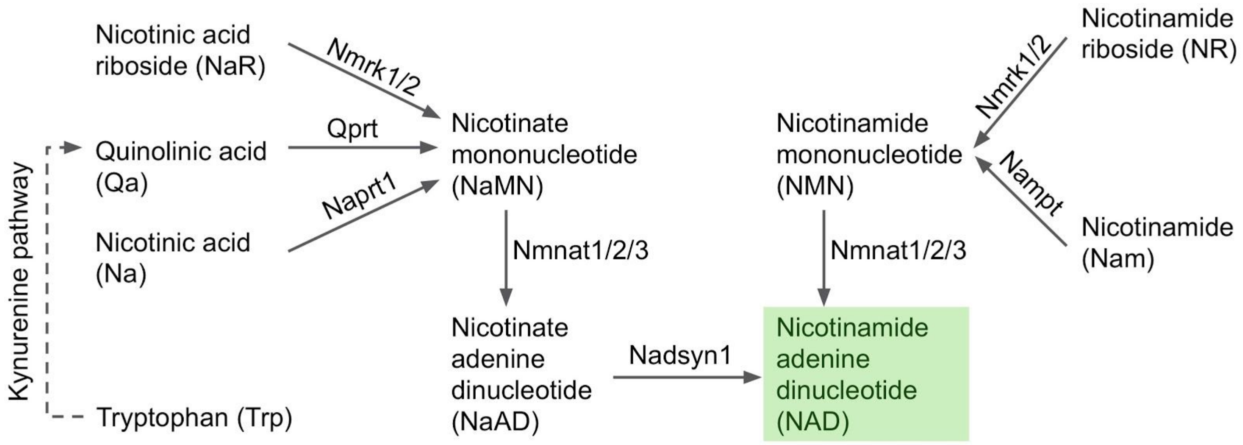


**Figure S5**. NAD biosynthesis pathway in mouse. The involved genes are Nmrk1 (nicotinamide riboside kinase 1), Nmrk2 (nicotinamide riboside kinase 2), Qprt (quinolinate phosphoribosyltransferase), Naprt1 (nicotinate phosphoribosyltransferase), Nmnat1 (nicotinamide nucleotide adenylyltransferase 1), Nmnat2 (nicotinamide nucleotide adenylyltransferase 2), Nmnat3 (nicotinamide nucleotide adenylyltransferase 3), Nadsyn1 (NAD synthetase 1), and Nampt (nicotinamide phosphoribosyltransferase).

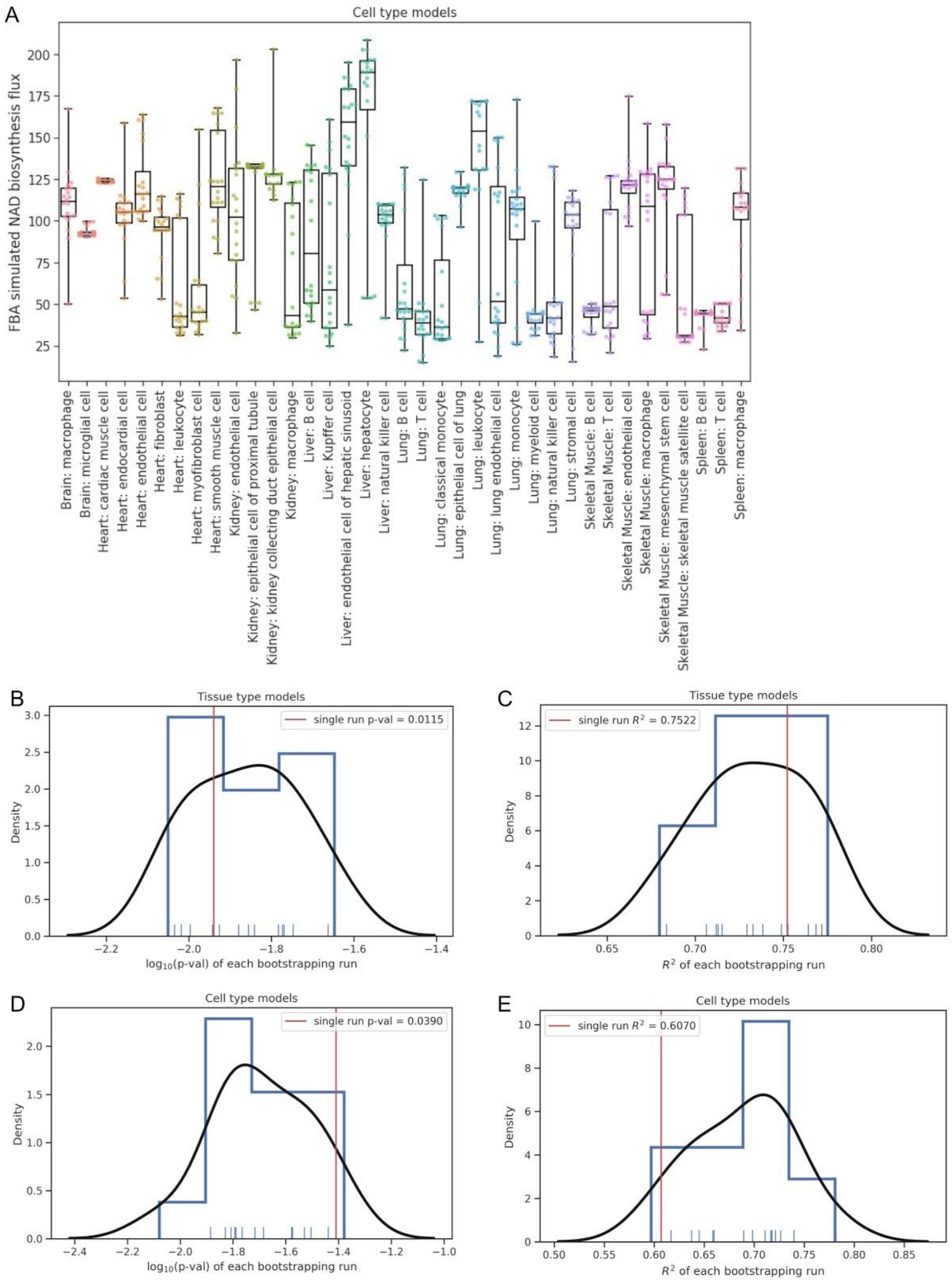


**Figure S6**. Bootstrapping results of metabolic modeling. (**A**) Distribution of the cell type model simulated $NAD^+$ biosynthesis fluxes in all bootstrapping samples. (**B, C**) Distribution of the p-values (**B**) and $R^2$ (**C**) of linear correlation between empirically measured $NAD^+$ fluxes and 15 bootstrapping runs of tissue type models. (**D, E**) Distribution of the p-values (**B**) and $R^2$ (**C**) of linear correlation between empirically measured $NAD^+$ fluxes and 15 bootstrapping runs of cell type models.

### Supplementary tables

**Table S1**. Linear correlations between empirically measured tissue specific $NAD^+$ biosynthesis fluxes and $log_2$ mean read counts + 1 of key enzymes in $NAD^+$ biosynthesis pathway.

| Gene symbol | R | $R^2$ | p-value | FDR |
|---|---|---|---|---|
| Qprt | 0.885 | 0.783 | 0.008 | 0.068 |
| Nmrk1 | 0.851 | 0.724 | 0.015 | 0.068 |
| Naprt1 | 0.659 | 0.434 | 0.107 | 0.322 |
| Nmnat1 | 0.596 | 0.355 | 0.158 | 0.356 |
| Nmnat2 | -0.279 | 0.078 | 0.545 | 0.645 |
| Nampt | 0.310 | 0.096 | 0.499 | 0.645 |
| Nadsyn1 | 0.380 | 0.144 | 0.401 | 0.645 |
| Nmrk2 | -0.260 | 0.068 | 0.573 | 0.645 |
| Nmnat3 | 0.109 | 0.012 | 0.816 | 0.816 |

**Table S2**. Simulated $NAD^+$ biosynthesis fluxes in different datasets. The mean and standard deviation (sd) flux values for $NAD^+$ biosynthesis over number of cells (n) simulated by 20 bootstrapping replicates at 80% of the cells per set. Cell types include astrocytes (Astro), endothelial cells (Endo), oligodendrocytes (Oligo) and neurons indicated by cortical layers 2-5 (L2-L5; L4 is not included in bootstrapping with only 3 cells). Intratelencephalic (IT), pyramidal tract (PT), and corticothalamic (CT). Lamp5: subclass of GABAergic neurons.

| Cell Type | ABI ALM NAD flux mean | ABI ALM NAD flux sd | ABI ALM cell n | ABI VISp NAD flux mean | ABI VISp NAD flux sd | ABI VISp cell n | TM NAD flux mean | TM NAD flux sd | TM n cells |
|---|---|---|---|---|---|---|---|---|---|
| Astro | 7.535 | 0.482 | 215 | 8.114 | 0.769 | 368 | 8.793 | 0.939 | 432 |
| Oligo | 9.876 | 1.665 | 98 | 10.450 | 1.765 | 91 | 11.844 | 1.665 | 1574 |
| Endo | 3.899 | 0.436148 | 715 | 4.828 | 2.997 | 94 | 4.578 | 0.436 | 715 |
| Macro | 9.690 | 1.594 | 85 | 10.550 | 1.579 | 51 | 11.031 | 2.143 | 61 |
| L2/3 IT | 13.086 | 1.168 | 325 | 14.053 | 1.547 | 982 | N/A | N/A | N/A |
| L5 IT | 7.793 | 0.000 | 2401 | 11.985 | 0.528 | 880 | N/A | N/A | N/A |
| L5 PT | 12.050 | 0.383 | 368 | 12.120 | 0.204 | 544 | N/A | N/A | N/A |
| L6 CT | 11.624 | 0.546 | 350 | 12.101 | 1.301 | 960 | N/A | N/A | N/A |
| L6 IT | 12.798 | 0.049 | 394 | 10.830 | 0.197 | 1872 | N/A | N/A | N/A |
| Lamp5 | 11.412 | 0.257 | 913 | 10.765 | 0.447 | 1122 | N/A | N/A | N/A |

#### Supplementary References


Brunk, E., Sahoo, S., Zielinski, D.C., Altunkaya, A., Dräger, A., Mih, N., Gatto, F., Nilsson, A., Preciat Gonzalez, G.A., Aurich, M.K., et al. (2018). Recon3D enables a three-dimensional view of gene variation in human metabolism. Nat. Biotechnol. *36*, 272–281.

Costa, R.S., Hartmann, A., and Vinga, S. (2016). Kinetic modeling of cell metabolism for microbial production. J. Biotechnol. *219*, 126–141.

Edwards, J.S., and Palsson, B.O. (2000). Robustness analysis of the escherichiacoli metabolic network. Biotechnol. Prog.

Fell, D.A. (2005). Enzymes, metabolites and fluxes. J. Exp. Bot. *56*, 267–272.

Gibson, B.A., and Kraus, W.L. (2012). New insights into the molecular and cellular functions of poly(ADP-ribose) and PARPs. Nat. Rev. Mol. Cell Biol. *13*, 411–424.

Hackett, S.R., Zanotelli, V.R.T., Xu, W., Goya, J., Park, J.O., Perlman, D.H., Gibney, P.A., Botstein, D., Storey, J.D., and Rabinowitz, J.D. (2016). Systems-level analysis of mechanisms regulating yeast metabolic flux. Science *354*.

Kosub, S. (2019). A note on the triangle inequality for the Jaccard distance. Pattern Recognit. Lett. *120*, 36–38.

Le Novère, N. (2015). Quantitative and logic modelling of molecular and gene networks. Nat. Rev. Genet. *16*, 146–158.

Megchelenbrink, W., Huynen, M., and Marchiori, E. (2014). optGpSampler: an improved tool for uniformly sampling the solution-space of genome-scale metabolic networks. PLoS One *9*, e86587.

Meindl, B., and Templ, M. (2012). Analysis of commercial and free and open source solvers for linear optimization problems. Eurostat and Statistics Netherlands within the Project ESSnet on Common Tools and Harmonised Methodology for SDC in the ESS *20*.

Orth, J.D., Thiele, I., and Palsson, B.Ø. (2010). What is flux balance analysis? Nat. Biotechnol. *28*, 245–248.

Palsson, B.O., and Lightfoot, E.N. (1984). Mathematical modelling of dynamics and control in metabolic networks. I. On Michaelis-Menten kinetics. J. Theor. Biol. *111*, 273–302.

Reid, M.A., Dai, Z., and Locasale, J.W. (2017). The impact of cellular metabolism on chromatin dynamics and epigenetics. Nat. Cell Biol. *19*, 1298–1306.

Saa, P.A., and Nielsen, L.K. (2017). Formulation, construction and analysis of kinetic models of metabolism: A review of modelling frameworks. Biotechnol. Adv. *35*, 981–1003.

Schaum, N., Karkanias, J., Neff, N.F., May, A.P., Quake, S.R., Wyss-Coray, T., Darmanis, S., Batson, J., Botvinnik, O., Chen, M.B., et al. (2018). Single-cell transcriptomics of 20 mouse organs creates a Tabula Muris. Nature.

Schultz, A., and Qutub, A.A. (2016). Reconstruction of Tissue-Specific Metabolic Networks Using CORDA. PLoS Comput. Biol. *12*, e1004808.

Sigurdsson, M.I., Jamshidi, N., Steingrimsson, E., Thiele, I., and Palsson, B.Ø. (2010). A detailed genome-wide reconstruction of mouse metabolism based on human Recon 1. BMC Syst. Biol. *4*, 140.

Thiele, I., and Palsson, B.Ø. (2010). A protocol for generating a high-quality genome-scale metabolic reconstruction. Nat. Protoc. *5*, 93–121.
